# Gait Adaptations Under Functional Asymmetry: Exploring the Role of Step Width, Step Length, and CoM in Lateral Stability

**DOI:** 10.1101/2024.12.23.630028

**Authors:** Tomislav Baček, Denny Oetomo, Ying Tan

## Abstract

Bipedal gait is inherently unstable, requiring a complex interplay between foot placement and centre of mass (CoM) movement to maintain balance. While various factors are known to impact walking balance, few studies have explored the specific effects of functional asymmetry on lateral stability. This study investigates how step length, step width, and CoM adaptations impact lateral gait stability in healthy young adults walking with and without a functional asymmetry induced by fully extending the left knee. The results show that step length remains unaffected by functional asymmetry regardless of speed, while step width increases under the constraint. This adjustment increases the base of support; however, the concurrent increase in lateral CoM movement reduces overall lateral stability. These findings offer valuable insights into fundamental gait adaptation and stability mechanisms, with potential implications for designing rehabilitation strategies for individuals with gait asymmetry.

## I. Introduction

The ability to maintain stability (i.e., avoid falling) is fundamental to human walking. This requires humans to keep their centre of mass (CoM) within a stable base of support (BoS) formed by the contact with the ground. Due to our CoM being located high above a relatively narrow BoS, bipedal gait is inherently unstable, necessitating multiple strategies to control balance during walking [1].

Precise foot placement has been identified as the primary mechanism for maintaining bipedal stability [2]. Foot placement control uses different mechanisms in the sagittal and frontal planes, but operates simultaneously and seemingly with a degree of interaction [2]–[4]. In the sagittal plane, anterior-posterior (AP) foot placement (i.e., step length, SL) primarily drives forward progression and contributes less directly to stability [4]. In contrast, medio-lateral (ML) stability in the frontal plane requires active control [3], which can be achieved either by modulating lateral foot placement (i.e., step width, SW) during the swing phase to adjust the next step’s BoS [5], [6] or by managing CoM during the stance phase [7].

While often studied separately, SW modulation and CoM movement are closely linked. Variations in CoM kinematics influence variations in SW [8], and the CoM trajectory during stance reliably predicts the next foot position variance [9]. This intricate interaction extends beyond lateral balance, as increased balance control – such as through enforced gait patterns – was shown to impose its own metabolic penalty [10], independent of costs associated with non-preferred SWs [11] or fluctuations in CoM energy [12]. Furthermore, the *bowtie* CoM trajectory, characteristic of healthy walking, changes uniformly with walking speed [13]. In contrast, SW changes with speed have shown conflicting trends: decreasing [14], remaining unchanged [15], following a U-shape [16], or exhibiting a reverse J-curve [17] as speed increases.

The interplay between SW and CoM dynamics becomes particularly relevant in the presence of functional asymmetry, which is common in neurologically impaired populations. Functional asymmetry often results in a disrupted *bowtie* CoM trajectory [18] and consistently wider SW across speeds [19]–[21], with falls, common during walking, often attributed to poor balance control [22]. However, studies in patient populations are often limited by confounding factors such as varying levels of impairment and individual-specific constraints, making it difficult to isolate the underlying biomechanical adaptations. Comparing patients to healthy controls matched by speed or age provides only partial insights, as inherent biomechanical and dynamic differences persist irrespective of these factors. A notable attempt to address this gap is McCain et al.’s study [23], which induced asymmetry in healthy individuals using knee and ankle bracing. However, the study was limited to walking only at 0.8 m/s, leaving functional asymmetry’s broader implications unexplored.

This study addresses these gaps by investigating gait adaptations under functional asymmetry in healthy young adults across three speeds, focusing on the interplay between SL, SW, and CoM dynamics. A unilateral knee constraint was used to emulate the challenges faced by individuals with impaired gait, such as stroke survivors, while avoiding the confounding factors present in patient studies. Stability was quantified using Hof et al.’s concept of the extrapolated CoM (xCoM) [24] and the margin of stability (MoS). We hypothesised that functional asymmetry would lead to significant adjustments in lateral and fore-aft foot placement to counter altered CoM dynamics. We also hypothesised that walking with a knee constraint would increase MoS to compensate for the heightened demands imposed by the asymmetry. This within-subject design provides new insights into biomechanical and stability strategies under functional asymmetry, with implications for both clinical and robotic gait interventions.

## II. Methodology

### A. Walking dataset

We used human gait dataset published by Baček et al. [25]. The dataset contains data of 21 neurotypical young adults who walked on a dual-belt instrumented treadmill over two sessions (days). In this study, data of only 17 participants are used since two dropped out after the first session (*Sub7* and *Sub12* in [25]) and two have poor lateral-medial ground reaction force (GRF) data (*Sub9* and *Sub10* in [25]). The 17 participants analysed herein include 5 females, with the following anthropometry: age 31 ± 8 years, body mass 72.8 ± 13.2 kg, height 1.71 ± 0.09 m (mean ± standard deviation).

All participants were free of any lower-extremity injury. Walking trials took five minutes each and include walking at three speeds (0.4, 0.8, and 1.1 m/s) and five cadences per speed (preferred, as well as two lower and two higher than preferred). All speed-cadence combinations were repeated twice: without constraints (i.e., *free*) and with a left knee joint extended using a passive knee brace (i.e., *constrained*), emulating hemiparetic compensatory gait. In this analysis, only tests at the participants’ preferred cadence from [25] are used: *T3, T8*, and *T13* for free walking, and *T18, T23*, and *28* for constrained walking at 0.4, 0.8, and 1.1 m/s, respectively.

### B. Gait variables

All gait variables were extracted and calculated in Matlab 2024a using custom-written scripts. GRF data were collected at 1 kHz and filtered using low-pass Butterworth filter with a 6 Hz cut-off frequency [26]. The vertical GRF component was used to segment walking data into gait cycles (time between two subsequent heel strikes of the same leg), with 5% of the peak amplitude used as a threshold (i.e., for a 70 kg person, the threshold is 70×9.81×0.05 = 34 N). All gait variables are an average over the last 60 seconds of a 5-minute trial, which ensured participants adapted to each walking condition before extracting data used to analyse their walking patterns [27].

We define step length (SL) as the fore-aft and step width (SW) as the medio-lateral distance between the two calcaneous (heel) markers at the time of the leading leg’s heel strike. Margin of stability (MoS) is calculated using the concept of extrapolated centre of mass (xCoM) [24]. Centre of mass (CoM) was calculated using the double integration method [28]; to calculate xCoM (1), CoM’s medio-lateral position (*pCoM*_*ml*_) was summed up <u>wi</u>th its corresponding velocity (*vCoM*_*ml*_) times a factor 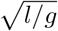, where *l* is the person’s leg length and *g* gravity acceleration. Medio-lateral MoS (2) was calculated as the smallest lateral distance between the centre of pressure’s (CoP’s) measured lateral position (*pCoP*_*ml*_) and the lateral xCoM position (*xCoM*_*ml*_) during the stance phase. Leg length was measured as the distance between the anterior superior iliac spine (ASIS) and the ipsilateral medial malleolus in a standing position during static calibration (see [25] for details). Positive MoS values indicate laterally stable gait (the higher the value, the more stable the gait is).

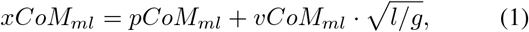

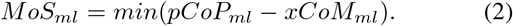

The medio-lateral CoM range of motion (ROM) is the distance between two furthest CoM positions in the lateral direction (frontal plane) during a gait cycle. A projection of the 3D CoM movement onto a transverse plane, i.e., the movement in the medio-lateral (ML) and anterior-posterior (AP) directions, makes a well-known shape resembling number eight [29]. We define the area enclosed by each of the number eight’s halves (*ellipses*) as CoM Area, and CoM Area Symmetry as a ratio between the left CoM Area divided by the sum of the left and right CoM Area (50% = symmetric).

### C. Statistical analysis

Statistical data analysis has been carried out in Python 3.10.10 using *scipy* package. The effects of speed and condition (free vs. constrained) on gait variables per leg were analysed using a two-way repeated measures ANOVA (RMANOVA) and a significance level of *p* = 0.05. Where statistically significant differences were found, Tukey’s Honestly Significant Difference (HSD) tests were conducted for pairwise post-hoc analysis, with Bonferroni corrections applied to account for multiple comparisons. Tukey’s HSD tests were also used to compare the difference between the two legs where applicable (Left vs. Right) for each speed-condition combination (e.g., free walking at 0.8 m/s). We define three levels of statistical significance: weak (0.01*<p*≤0.05), moderate (0.001*<p*≤0.01), and strong (*p*≤0.001).

### III. Results

In all bar graphs, three walking speeds are separated by vertical dashed lines, and two conditions (free vs. constrained) are colour-coded in blue and red, respectively. Data are average across all 17 participants, with full bars representing mean and error bars standard deviation of the population data.

### A. Foot placement – step length (SL) and width (SW)

Increasing gait speed increases SL on both legs (Fig. 1, top row), with statistically significant effect of speed (strong effect, *p<*0.001) during both free (blue bars) and constrained (red bars) walking. SL symmetry (not visualised), calculated as the ratio of the left SL to the sum of the two SLs (50%=symmetric), remains consistent and strongly symmetric – free vs. constrained, on average: 50.2% vs. 49.3% at 0.4 m/s, 50.1% vs. 51.1% at 0.8 m/s, and 49.9% vs. 50.4% at 1.1 m/s. Consequently, across all speeds and conditions, there are no statistically significant differences in SL symmetry, or when comparing free to constrained SL (*p>*0.85).

**Figure 1.**
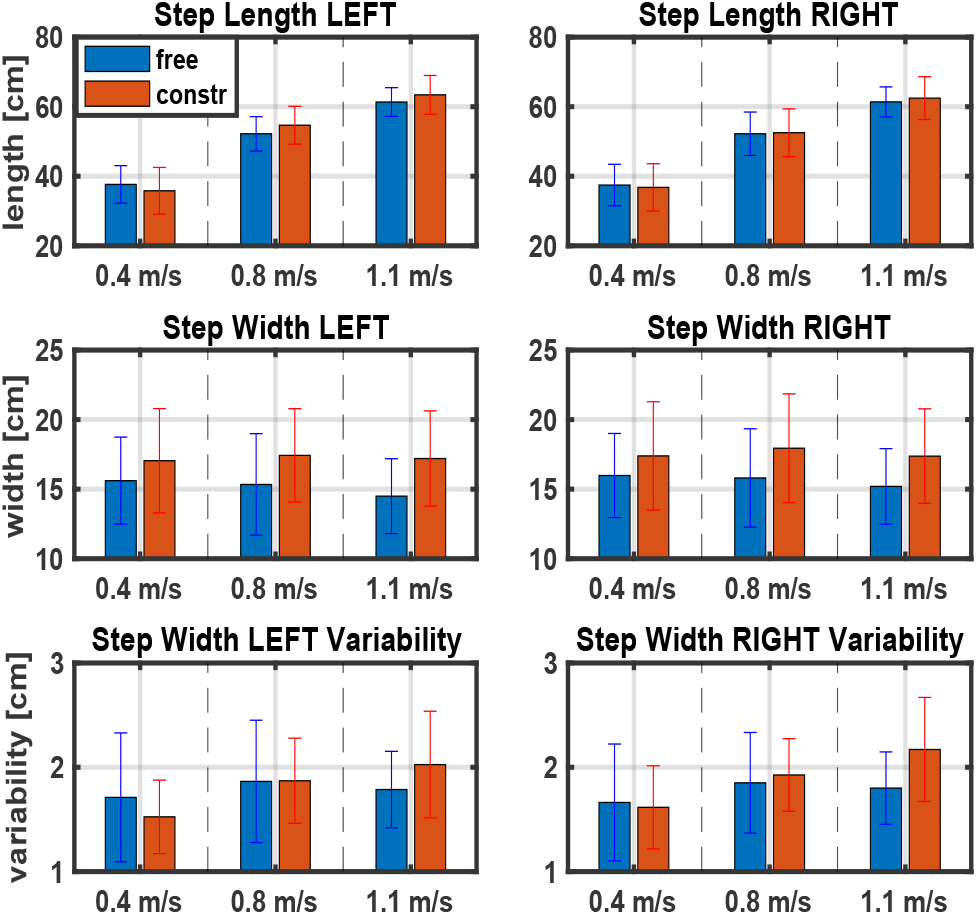
Step length (SL), Step width (SW), and SW variability across conditions. Note the same y-axis in each row. SW variability is calculated by averaging individual variability across participants.

Increasing walking speed has little effect on SW irrespective of the walking condition (Fig. 1, middle row); on both legs and in both conditions (free, constrained), the effect of speed is statistically insignificant (*p>*0.95). Adding a passive knee brace increases SW by about 2 cm (≈12%) on both legs, but the increase is not statistically significant (*p>*0.12). Similarly, there are no differences in SW between Left and Right legs in any of the speed-constraint combinations (*p>*0.99).

A different trend is observed in SW variability (Fig. 1, bottom row). During free walking, there is no statistically significant effect of speed (*p>*0.49) or side (Left vs. Right; *p>*0.99) on repeatability of lateral foot positioning. When a knee constraint it added, walking speed only has a statistically significant effect when comparing 1.1 to 0.4 m/s on both legs (moderate effect, 0.001*<p<*0.01). SW variability remains similar across the two legs (*p>*0.93), and added constraint does not statistically significantly change it on either leg.

Overall, knee constraint does not affect SL or SL symmetry, indicating a disassociation from SW modulation. SW is unaffected by the speed but increases during constrained walking, suggesting participants used SW to control lateral stability.

### B. Margin of stability (MoS)

Increasing walking speed increases lateral MoS in both free (Fig. 2, top row, blue bars) and constrained walking (Fig. 2, top row, red bars), albeit without statistical significance (*p>*0.21). Similarly, no statistically significant differences exist when comparing MoS between the two legs at any of the six speed-constraint combinations (*p>*0.59). Adding a knee brace on the left leg reduces average MoS on both legs and across all three speeds (Fig. 2, top row, blue vs. red) albeit with no statistical significance (*p>*0.17). The exception is the left leg at 0.8 m/s, which exhibited a decrease as big as 35% (or 3 cm; *p*=0.07). Note that, on average, participants always walk with a stable gait (MoS*>*0); individually, two participants had MoS*<*0 on their left leg during free walking at 0.4 m/s; during constrained walking, the same two participants had MoS*<*0 on their left leg at 0.4 and 0.8 m/s, and one more participant at 0.4 m/s.

**Figure 2.**
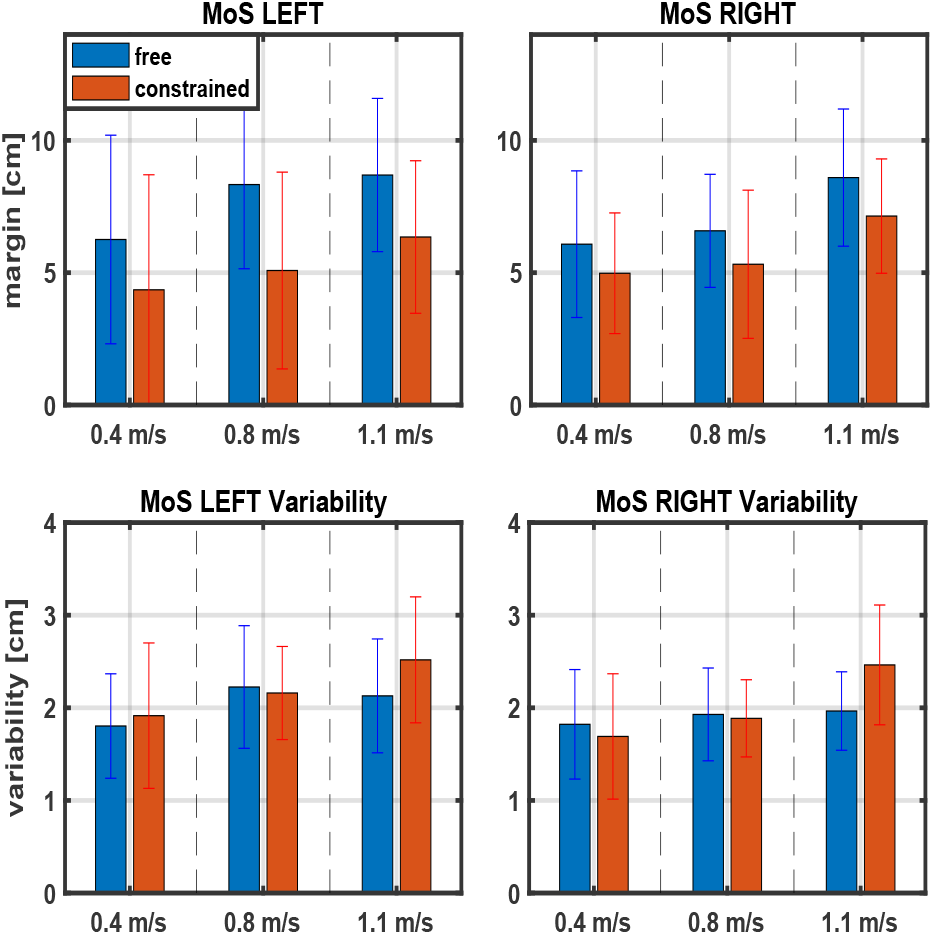
Medio-lateral margin of stability (MoS) and its variability across conditions. Note the same y-axis limits in each row. Positive values indicate stable gait.

MoS variability remained fairly consistent across speeds, conditions, and legs (Fig. 2, bottom row). There is no statistically significant effect of speed in either leg during free walking (*p>*0.28); during constrained walking, speed only statistically significantly changes MoS variability when comparing 1.1 to 0.4 m/s on the left (weak effect, *p*=0.04) and right leg (strong effect, *p*=0.0007). There are no Left vs. Right differences in either free or constrained walking (*p>*0.54), and adding constraint to the left knee introduces no statistically significant changes to MoS variability in either leg (*p>*0.42; notably, *p*=0.06 on the right leg at 1.1 m/s).

Overall, MoS slightly increased with speed but decreased on both legs under knee constraint, though without statistical significance. Trends in MoS variability suggest that participants found it more challenging to control MoS during constrained walking only at 1.1 m/s, and equally so on both legs.

### C. Centre of Mass (CoM) movement

As walking speed increases, lateral CoM movement tends to decrease statistically significantly in both free and constrained walking (strong effect, *p<*0.001; Fig. 3, top left). Notably, despite an increase in the lateral CoM movement when walking with constraints, the change is not statistically significant at any speed compared to free walking (*p>*0.67). In contrast, CoM area ratio notably changes (Fig. 3, top right). During free walking (blue bars), participants maintained near-perfect symmetry across speeds (*p>*0.98), as illustrated by the CoM path in the transverse plane (Fig. 3, bottom, full lines). Under constraints, CoM area symmetry dropped below 50%, indicating greater reliance on the right (unconstrained) leg. The effect of speed on CoM area ratio during constrained walking leads to statistically significant difference between 0.4 and 1.1 m/s (strong effect, *p*=0.0003), and 0.4 and 0.8 m/s (moderate effect, *p*=0.008), and between free and constrained walking at 0.8 and 1.1 m/s (strong effect, *p<*0.001).

**Figure 3.**
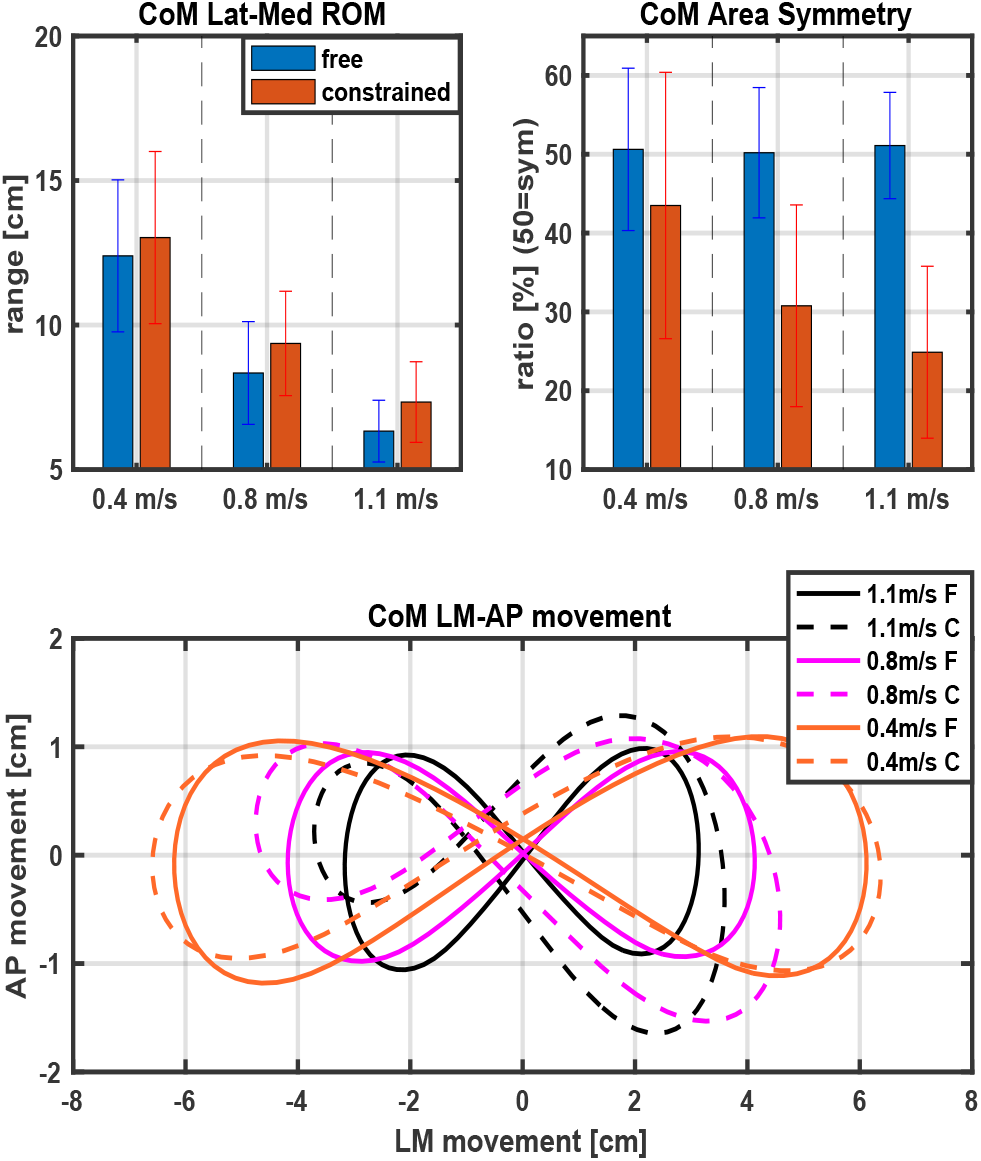
Centre of mass (CoM) movement in the frontal plane (ROM) and projected onto the transverse plane (Area Ratio, LM-AP movement). Free and constrained conditions are colour-coded in the top row; in the bottom row, walking speeds are colour-coded and the two conditions visualised using full (free; F) and dashed (constrained; C) lines.

Dashed lines in Fig. 3 (bottom) illustrate the impact of a knee brace on CoM trajectory. At 0.8 and 1.1 m/s, functional asymmetry increases both lateral and fore-aft movement of the CoM on the right side, while only lateral movement increases on the left side. At 0.4 m/s, lateral CoM movement increases slightly on the right side, with reduced fore-aft and increased lateral movement on the left side. The constraint-induced changes are less pronounced at 0.4 m/s, explaining the smaller free vs. constrained CoM area ratio difference at this speed compared to the two faster speeds (Fig 3, top right).

Overall, lateral CoM movement decreases with speed during both free and constrained walking, with the constraint having the greatest impact in the fore-aft direction. Constrained walk-ing also results in marked asymmetry in the CoM area ratio, reflecting a compensatory reliance on the right (unconstrained) leg to control lateral stability.

## IV. Discussion

This study investigated human gait adaptations under functional asymmetry across a range of speeds relevant to healthy young adults (1.1 m/s, [30]) and hemiparetic patients (0.8 and 0.4 m/s, [31]). By having participants serve as their own control, we isolated these adaptations from confounding factors often present in patients or between-group comparisons. The primary focus was on the interplay between step length (SL), step width (SW), and centre of mass (CoM) movement and their combined effects on lateral gait stability. Our first hypothesis was partially confirmed: participants modulated SW but not SL to counteract CoM impacts on lateral stability. Conversely, our second hypothesis was rejected, as MoS decreased, rather than increased, during constrained walking.

### A. Walking dataset

Human gait adaptations are often studied by comparing different groups, such as through speed-or age-matching designs [32], [33]. However, these approaches cannot fully account for biomechanical differences that are independent of speed or age, especially in patient populations [34]. The within-subject design of the walking study used herein [25], where participants act as their own control, offers a unique advantage by isolating gait adaptations solely due to functional asymmetry, free from confounding factors (e.g., muscle weakness). In the study, participants wore a passive knee brace that restricted left knee flexion, limiting ankle plantarflexion during push-off and inducing propulsive asymmetry [35] as well as compensatory gait adjustments [36]. As gait speed increased, propulsive asymmetry did too [35], reflecting the natural demands of higher speeds on joint mobility [37]. Despite a metronome guidance, participants were free to make slight adjustments to their step length and step time (and their symmetries) as long as the resulting gait speed (i.e., *stride* length / *stride* time) matched that imposed by the treadmill.

### B. Fore-aft foot placement

During free walking, participants increased their SL with speed (Fig.1, top row, blue bars), consistent with the well-documented positive correlation between gait speed and SL [38]. This adaptation was observed symmetrically across both legs, resulting in strong SL symmetry at all three speeds, concurrent with the speed-agnostic propulsive symmetry reported by [35]. These findings reinforce the critical role of SL in forward progression and its assumed influence on the mechanical and metabolic costs of walking [11], [39].

Interestingly, introducing a left knee constraint had no measurable effect on SL or its symmetry (Fig.1, top row, red bars), even in the presence of pronounced propulsive asymmetries at 0.8 and 1.1 m/s [35] (similar to those observed in stroke patients classified as having *mild* to *moderate* impairments [40]). These results align with findings by McCain et al. [23], who similarly used knee brace to externally induce propulsive asymmetry and reported minimal impact on SL symmetry (and metabolic cost) during walking at 0.8 m/s. Importantly, this consistency holds despite differences in our experimental protocols, as McCain et al. did not employ metronome guidance.

A notable observation is that their participants prioritised SL over step time (ST) symmetry during constrained walking. Although ST symmetry is beyond the scope of this paper, we note that our participants exhibited similar adaptations. This behavior contrasts with findings from free walking, where ST, rather than SL asymmetry, has been shown to significantly influence the metabolic cost of walking [12], [41]. The underlying mechanisms driving this prioritisation during constrained walking remain unclear and warrant further investigation.

### C. Lateral foot placement and medio-lateral stability

In the walking study used herein [25], participants on average maintained a consistent SW and SW variability across different speeds during free walking, with similar values observed for both legs (Fig. 1, middle and bottom rows, blue bars). This stability in SW across speeds aligns with findings from [6] and [15], though it contrasts with studies like [14], [16], [17], likely due to differences in experimental designs and speed ranges tested. Additionally, our results support the finding in [4] of an independent SW-SL control at self-selected speeds, extending it across a wider range of speeds.

Generally, higher walking speeds during free walking result in reduced CoM movement in 3D space [29], as there is less time to execute step-to-step transitions. When this decrease is decoupled from SW modulations, as was the case here – SW remained consistent across speeds (Fig. 1, middle row, blue bars) – lateral stability, as defined by MoS, would be expected to increase. As shown in Fig. 2, this was indeed the case. Using MoS at 1.1 m/s, a speed close to the preferred metabolically efficient gait speed for healthy young adults, as a reference for stability, the reduction in MoS at 0.4 m/s, despite participants reporting feeling *less stable* and disliking the speed, suggests that lateral stability may not be prioritised at slower speeds.

This is likely due to the metabolic penalty associated with wider steps. While taking wider steps at a given speed enhances lateral stability in both healthy individuals and hemiparetic patients [32], [33], [42], it also increases metabolic cost [11]. Furthermore, wider steps are typically more constrained at higher speeds, where longer steps are preferred [42], suggesting that participants could have taken wider steps at lower speeds to increase their MoS. The fact that they didn’t – in fact, quite the opposite – suggests that a combination of having more time to correct for potential disturbances at slower speeds and the intrinsic metabolic penalty of walking slowly takes precedence over maximising lateral stability, which is instead maintained at a *sufficient* level during free walking.

Adding a knee constraint impacted SW, MoS, and CoM, but to different degrees. Walking with propulsive asymmetry, particularly at 0.8 and 1.1 m/s [35], led to significant asymmetry in CoM area (Fig. 3, top right), indicating increased reliance on the right (unconstrained) leg. The fully extended left knee prevented participants from performing *natural* weight acceptance following heel strike or push-off after toe-off, resulting in reduced CoM movement in the AP and increased in the LM directions (Fig. 3). The apparent effect of increased lateral CoM movement at a given speed is comparable to reducing speed, which typically allows more time to manage potential disturbances. This increased need for lateral stabilisation was accompanied by an increase in SW across speeds, as expected, and is consistent with the literature [7], [33], [43].

The increase in SW during constrained walking was consistent across speeds, mirroring the trend observed during free walking. This could be specific to our group of participants, whose preference for a consistent SW across conditions may not have been significantly challenged by the constraint or the range of tested speeds. As a result, they may have defaulted to habitual motor patterns developed during free walking, potentially trading MoS at slower walking speeds. However, this does not fully explain why participants walked with decreased MoS across all speeds compared to free walking.

At 1.1 m/s, it’s conceivable that their SW was near its max value, making it mechanically difficult to walk wider. This hypothesis is supported by the increased variability in SW and MoS compared to other conditions (Fig. 1 and 2). At 0.8 and 0.4 m/s, however, this explanation seems unlikely. Instead, altered CoM dynamics during constrained walking (Fig. 3), also common in neurologically-impaired patients [18], combined with the metabolic penalties associated with increased SW and lateral stabilisation, suggests that the trade-off landscape between lateral stability and metabolic cost was shifted by the constraint. If true, participants may still have been minimising their metabolic cost, even at the expense of reducing their stability margin. This hypothesis, though, is beyond the scope of this paper and warrants further investigation.

### D. Limitations

This study has several limitations. First, the participant pool of young, healthy adults limits the generalisability of the findings to broader populations. Second, it focused on SL, SW, and CoM dynamics, excluding variables like step time symmetry and metabolic cost. Lastly, using MoS as the sole metric for lateral stability, while valuable, may not fully capture the complexity of dynamic balance recovery strategies.

## V. Conclusion

This paper examines gait adaptations in young, healthy adults walking with and without propulsive asymmetry, focusing on how step length, step width, and lateral centre of mass (CoM) movement interact to influence lateral margin of stability. Results indicate that humans prioritise step length symmetry even under propulsive asymmetry, while maintaining consistent step width across speeds. This results in a slight increase in lateral stability with higher speeds, suggesting humans prioritise a trade-off that ensures *sufficient* gait stability. Functional asymmetry significantly changes gait dynamics, including CoM movement and stability margins. These adaptations highlight a complex interplay between lateral balance, preferred motor patterns, and potentially metabolic efficiency. Overall, this underscores the importance of targeting propulsive asymmetry and CoM dynamics in correcting gait, rather than focusing on, e.g. reducing step length asymmetry.

## Acknowledgment

The authors would like to thank Mr. Yufan Xu and his supervisor, Dr. Liuhua Peng of The University of Melbourne’s Faculty of Science, for providing their expertise in running statistical analysis and preparing results.

